# Analysis of *Plasmodium vivax* schizont transcriptomes from field isolates reveals heterogeneity of expression of genes involved in host-parasite interactions

**DOI:** 10.1101/2020.03.20.998294

**Authors:** Sasha V. Siegel, Lia Chappell, Jessica B. Hostetler, Chanaki Amaratunga, Seila Suon, Ulrike Böhme, Matthew Berriman, Rick M. Fairhurst, Julian C. Rayner

## Abstract

*Plasmodium vivax* gene regulation remains difficult to study due to the lack of a robust *in vitro* culture method, low parasite densities in peripheral circulation and asynchronous parasite development. We adapted an RNA-seq protocol “DAFT-seq” to sequence the transcriptome of four *P. vivax* field isolates that were cultured for a short period *ex vivo* before using a density gradient for schizont enrichment. Transcription was detected from 78% of the PvP01 reference genome, despite being schizont-enriched samples. This extensive data was used to define thousands of 5’ and 3’ untranslated regions (UTRs), some of which overlapped with neighbouring transcripts, and to improve the gene models of 352 genes, including identifying 20 novel gene transcripts. This dataset has also significantly increased the known amount of heterogeneity between *P. vivax* schizont transcriptomes from individual patients. The majority of genes found to be differentially expressed between the isolates lack *Plasmodium falciparum* homologs and are predicted to be involved in host-parasite interactions, with an enrichment in reticulocyte binding proteins, merozoite surface proteins and exported proteins with unknown function. An improved understanding of the diversity within *P. vivax* transcriptomes will be essential for the prioritisation of novel vaccine targets.

## Introduction

*Plasmodium vivax* is the second most prevalent malarial infection worldwide, and infection rates are continuing to increase as many global elimination strategies remain focused on *P. falciparum* malaria^1,2^. *P. vivax* is the dominant malaria species in Southeast Asia, South America, and Northeast Africa. Advances in the biological understanding of *P. vivax* are inhibited by the lack of a continuous culturing method and a corresponding shortage of functional assays in comparison to those available for *P. falciparum*, which has been broadly studied after culture adaptation was first achieved in 1976^3^. Since the absence of continuous *in vitro* culturing methods restrict experimental progress, most techniques to study *P. vivax* rely on parasites sampled directly from patient donors, but these are limited in quantity and can be challenging to access. Studies relying on *P. vivax* clinical parasite samples have additional challenges to overcome because of the mixture of parasite blood stages that are asynchronous in peripheral circulation (due to the lack of sequestration in this species), as well as frequent polyclonal infections and difficulties obtaining sufficient parasite genetic material for downstream analysis ^4–7^.

The first reference genome available for *P. vivax* was the Salvador-I (Sal1) reference genome, which is highly fragmented (>2500 unassembled scaffolds). *P. vivax* genomes have substantial geographically-distinct genetic diversity and a significant proportion of the disease burden lies in Southeast Asia, whereas the Sal-I strain was originally isolated in El Salvador^8^. Recently, a new reference genome from a Papua Indonesian isolate was released that combined short- and long-read sequencing technology, and consequently made considerable improvements in assembly and annotation (PvP01) compared to Sal1. This has greatly improved the understanding of *P. vivax* genome structure and read mapping for patient isolates^9^.

The first transcriptomic studies of *P. vivax* from field isolates and its comparison with other *Plasmodium* species were performed with microarrays, and showed similar cascades of stage-specific gene expression found in other *Plasmodium* species, where most genes show a clear peak of transcript at a specific time point in the intraerthyrocytic developmental cycle (IDC), with regulation presumed to involve ApiAP2 DNA-binding proteins ^10–13^. The first study to use RNA-seq in *P. vivax* was able to describe the features of the transcriptome more comprehensively, as previously unannotated untranslated regions (UTRs) and splice sites were outside the technical capabilities of microarrays^14^. Other previous studies of bulk RNA-seq from patient isolates have been performed on both a time course and with asynchronous samples^15–17^, and with primates in laboratory conditions ^18^.

Here we describe RNA-seq data from purified schizonts collected from four Cambodian patient isolates that had been subject to short-term *ex vivo* culture to allow for parasite maturation. The data was generated with minimal PCR amplification, based on the DAFT-seq protocol (directional, amplification-free transcriptome sequencing); this data was able to help further refine and increase the resolution of the annotation of the PvP01 reference genome, including identification of novel transcripts and correction of several hundred gene models. The increased evenness in coverage in this dataset provides an improved view of the *ex vivo* schizont transcriptome, as a large number of PCR amplification cycles in a standard RNA-seq library preparation induces bias by preferentially enriching for GC-rich regions of the genome, and reducing coverage in the most AT-rich regions where UTRs and ncRNAs are present. We also describe new features of the transcriptome, such as transcription start site associated RNAs (TSSa-RNAs) and intron-like features overlapping protein-coding exons (exitrons). We also generated a comprehensive list of enriched *P. vivax* schizont genes which provides the potential of a more in-depth understanding of *P. vivax* merozoite invasion, which to date is largely limited to a comparison with *P. falciparum* invasion homologs and a handful of known *P. vivax* invasion gene families such as the duffy binding like proteins (DBLs), reticulocyte binding proteins (RBPs), and merozoite surface proteins (MSPs)^19–22^.

## Results

### Preparation of purified late-stage schizont transcriptomes

Four blood samples from Cambodian patients were selected to undergo short-term *ex-vivo* culture. After the majority of parasites had matured, as judged by microscopy, late-stage schizonts were purified using Percoll gradients. Four RNA-seq libraries were generated using a modified version of the DAFT-seq protocol, which was optimised for highly AT-rich *Plasmodium* parasites^23^, and sequenced using the Illumina platform to generate 55-63 million reads per patient sample. In all cases >85% of reads mapped to the *P. vivax* PvP01 reference genome (**Table S1**). This data was used to improve the gene models of 352 genes in the PvP01 genome, including identifying 20 novel gene transcripts (Table S2). Comparison of the expression values of these RNA-seq libraries to blood stage microarray time course from a *P. vivax* dataset (containing three patient isolates)^12^ **(Fig. S1)** and a more densely sampled *P. falciparum* dataset^24^ **(Fig. S2)** confirmed that the samples were late-stage schizonts (as they correlated most strongly to these time points in the prior datasets) and that the transcriptomes were highly similar to each other (as the correlation of the normalised expression values of the RNA-seq libraries was close to 1) **(Fig. S3)**. More apparent heterogeneity was found between the patient isolates using the PVP01 genome than using only the gene IDs present in the Sal1 genome **(Fig. S4)**, consistent with most of the heterogeneity within the patient isolate transcriptomes being present in multigene families that are difficult to assemble and annotate, and are thus under-represented in the Sal1 genome.

### The architecture of the *Plasmodium vivax* schizont transcriptome is dense and overlapping

We used the RNA-seq data from the four patient isolates to define the extent of the *P. vivax* late-stage schizont transcriptome. We were able to detect transcription from 78% of the genome sequence using a threshold of 5 reads (as used in Siegel *et al*. 2018^25^). We were able to detect expression of 5,017 protein-coding genes (75% of annotated genes) at a threshold of 5 reads per kilobase per million mapped reads (RPKM), with the majority (3,974 genes, 60% of annotated genes) of these genes also detected as expressed at a much more conservative threshold of 20 RPKM **(Table S3)**.

We defined the 5’ untranslated regions (UTRs) for 4,155 genes in the PvP01 genome using the RNA-seq data (**Fig. 1a**, **Table S4**) (83% of those detected at >5 RPKM). We also defined 3’ UTRs for 4,091 genes (**Fig. 1b**, **Table S5**) (82% of those detected at >5 RPKM). We used an approach developed for *P. falciparum* RNA-seq data^23^ to define UTRs in AT-rich regions. This computational approach **(Fig. S5)** relies on RNA-seq data mapping continuously across the length of an mRNA, which is a key strength of the RNA-seq protocol used in this study. The precise boundary of the UTR for each molecule is likely to vary slightly, so to annotate a single fixed position for each mRNA boundary, we estimated a true position by defining the location where the continuous RNA-seq coverage falls below a threshold of 5 reads. To avoid merging adjacent UTRs on the same strand we used a more stringent threshold; the threshold for defining a block of continuous transcription was iteratively increased, in steps of 5 reads. The mean length of 5’ UTRs was 1,007 nt (median length 815 nt), and was 818 nt for 3’ UTRs (median was 685 nt), which is longer than those described in previous studies in *P. vivax*^*14,15*^. These are also longer than the 5’ and 3’ UTR lengths recently described in *P. falciparum*^*23*^ (577 nt and 453 nt, respectively). This difference is likely primarily technical in nature, due to many fewer sequences of >90% AT content disrupting continuous coverage in the *P. vivax* transcriptome relative to the *P. falciparum* transcriptome. The longest 5’ UTR detected was 6,846 nt, which belonged to the gene AP2-G (PVP01_1440800) **(Fig. S6)**, which also had the longest 3’ UTR (5,908 nt). Together these extended UTRs allow us to define that the majority of the genome (at least 73%) encodes mRNAs; this is an underestimate of the true value, as not all genes are expressed in these samples which are highly enriched for schizonts.

**Figure 1:**
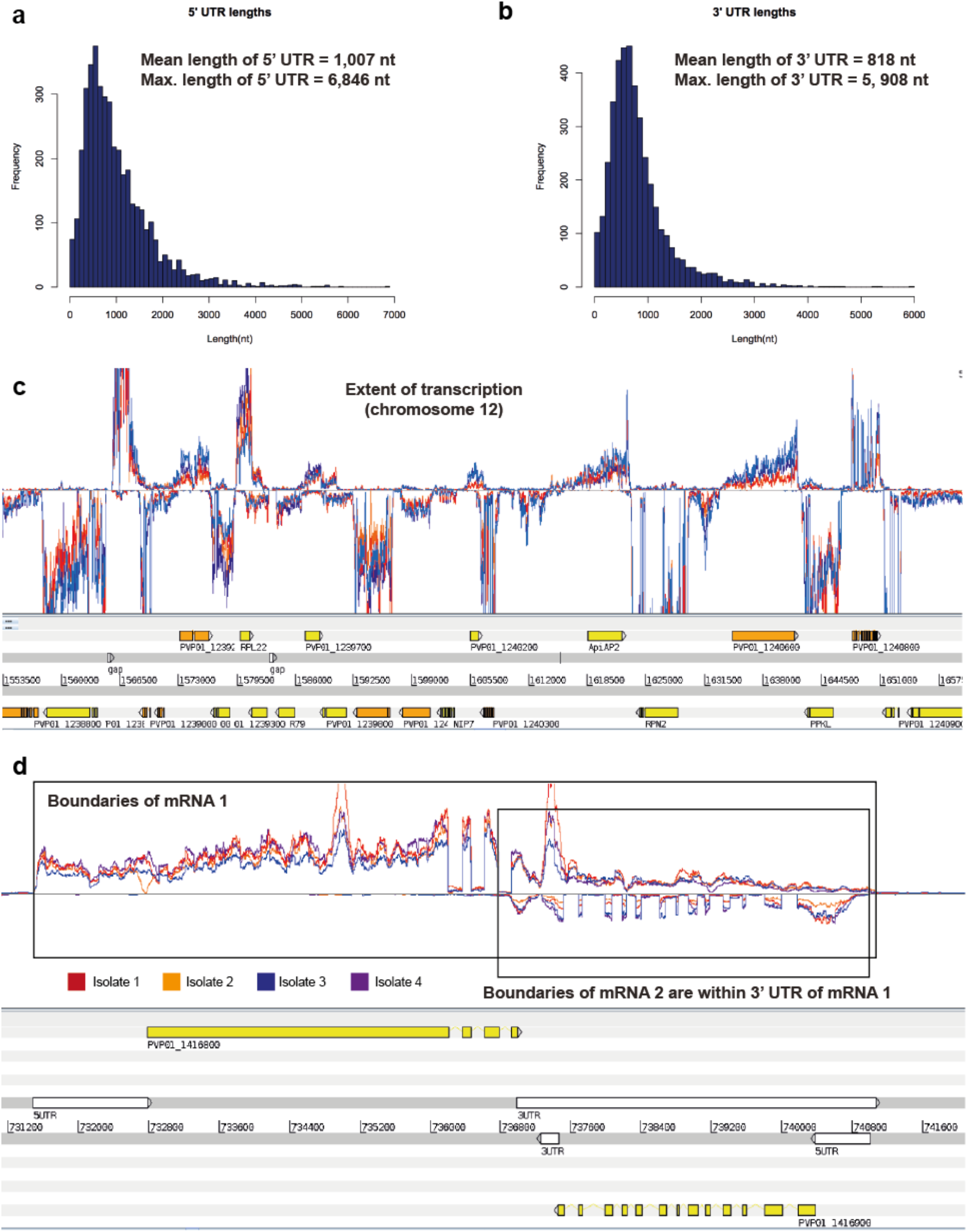
The extent of the *P. vivax* schizont transcriptome. a. Size distribution of 5’ UTRs (n =4155) b. Size distribution of 3’ UTRs (n=4,091) c. An overview of the extent of transcription from a representative portion of the *P. vivax* genome sequence (from chromosome 12). The coloured lines in the upper panel represent directional RNA-seq coverage from each of the four patient isolates, while the lower panel includes gene models on both strands of the genome sequence. d. Overlapping transcripts are found even within a single life stage in *P. vivax*. The example shown is of a gene pair in a “tail-to-tail” orientation (PVP01_1416800 and PVP01_1416900). The boundaries of the mRNA sequence of the second gene in this pair is contained within the 3’ UTR sequence of the first gene in the pair.

A region of the genome containing multiple genes detected in schizonts is shown in **Fig. 1c**. A significant number of genes substantially overlap each other, and we detected hundreds of overlapping transcripts even within this specific life stage. There were 1822 genes where the 3’ UTRs overlapped. An example of a gene pair in a “tail-to-tail” orientation (where the genes are on opposite strands, PVP01_1416800 and PVP01_1416900) is shown in **Fig. 1d.** The boundaries of the mRNA sequence of the second gene in this pair is contained within the 3’ UTR sequence of the first gene in the pair. We found many examples of genes in a “head-to-head” orientation that appear to share a bidirectional promoter. In some cases these 5’ UTRs are directly adjacent (**Fig. S7a)**, while others (968 genes) show some overlap (**Fig. S7b)**.

### Identification of new splice sites and non-coding transcription

We also used the RNA-seq data to search for additional non-coding transcripts, using blocks of continuous coverage like those used for detecting UTRs. We found evidence of non-coding transcripts showing spatial patterns of expression similar to the patterns of transcription start site associated RNAs (TSS-associated RNAs) recently described in *P. falciparum* ^23^, which are found in an antisense orientation to the 5’ end of mRNAs, such as the example shown upstream of the gene AP2-G3 (PVP01_1418100) (**Fig. S8)**.

We were also able to use the RNA-seq data to detect splice sites. We used an approach previously applied to *P. falciparum* DAFT-seq data^23^, where all spliced reads are examined and categorised. We found 8,217 splice sites that were already annotated for the PvP01 genome available in online databases, to which we added 7,418 new splice sites (**Fig. 2a**, **Table S6)**. We found 3,015 splice sites in UTRs, including 282 that affect the position of start or stop codons, as well as 2,164 splice sites enabling isoform variation within protein-coding exons. We also found evidence for alternative splicing in 2.3 % of coding genes. This proportion is lower than estimates in *P. falciparum* DAFT-seq data^23^, but this data set covers only a portion of the IDC.

**Figure 2:**
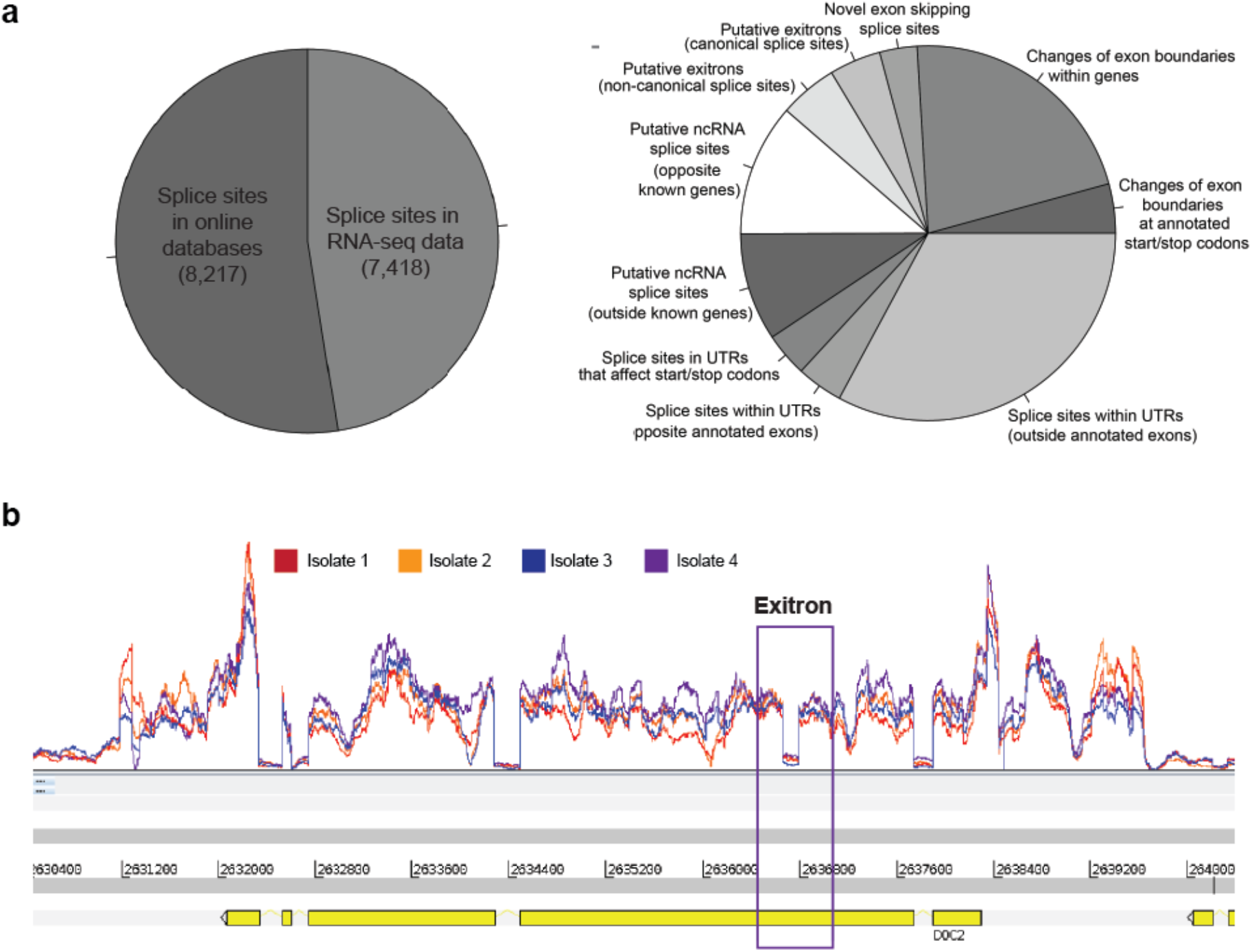
Splice sites present in *P. vivax* schizonts. a. Thousands of splice sites were detected in the RNA-seq data (left), with splice sites not found in online databases (right) falling into a range of categories for both coding and non-coding regions of RNAs. b. An exitron was identified in RNA-seq data for the gene PVP01_1461000. This exitron is 132 nt, a multiple of 3 nt, which can be spliced out without changing the reading frame of this protein. The vast majority of the transcripts for this mRNA contain the spliced form of the exitron.

In addition to conventional splice sites, we also found evidence for the presence of exitrons in *P. vivax* transcripts, which are intron-like features within exons where splicing can change protein sequence and hence increase proteome diversity^26^. Crucially, exitron splicing does not necessarily maintain the open reading frame of the protein, causing a diversification of protein sequences. An equally important role may be enabling another layer of regulation of transcript levels through mechanisms such as nonsense-mediated decay of transcripts that would produce non-functional proteins. Putative exitrons have recently been identified in P. falciparum^23^. Using our pipeline, there were 702 splicing events within exons that show some exitron properties (**Table S7**). For example, for PVP01_1461000, 132 nt (a multiple of 3 that retains the open-reading frame) is spliced out in the vast majority of the detected transcripts (**Fig. 2b**). Detailed mechanistic studies would be needed to establish the functions of these exitrons in *Plasmodium* species.

### Identification of patterns of similarity and heterogeneity between patient isolates

As part of the sample generation process, Percoll-purification was used to isolate schizonts, which has the advantage of eliminating the mixture of asexual blood stages typically seen in clinical infections. As a result we could use this data to characterise heterogeneity between *P. vivax* schizont transcriptomes. We first looked at the genes with the highest abundance in each patient isolate and found that there was a large overlap in the highest schizont-expressed genes in each sample, with 20 of the top 25 most highly expressed genes being present among the top 25 most highly expressed genes for all four patient isolates **(Table 1).** Several of the highest expressed genes belong to families implicated in host-parasite interactions, such as the merozoite surface proteins (MSPs), and early transcribed membrane proteins (ETRAMPs), with the majority of highest expressed genes being those that encode proteins involved in translation. Both histones and ribosomal RNA subunits were highly expressed, and expected to be seen in mature parasite stages undergoing active replication. Patterns of RNA-seq coverage across the genome are also highly conserved among patient isolates, with chromosome 12 as a representative example showing the vast majority of genes being transcribed at very similar levels (**Fig. 1c**).

**Table 1.**
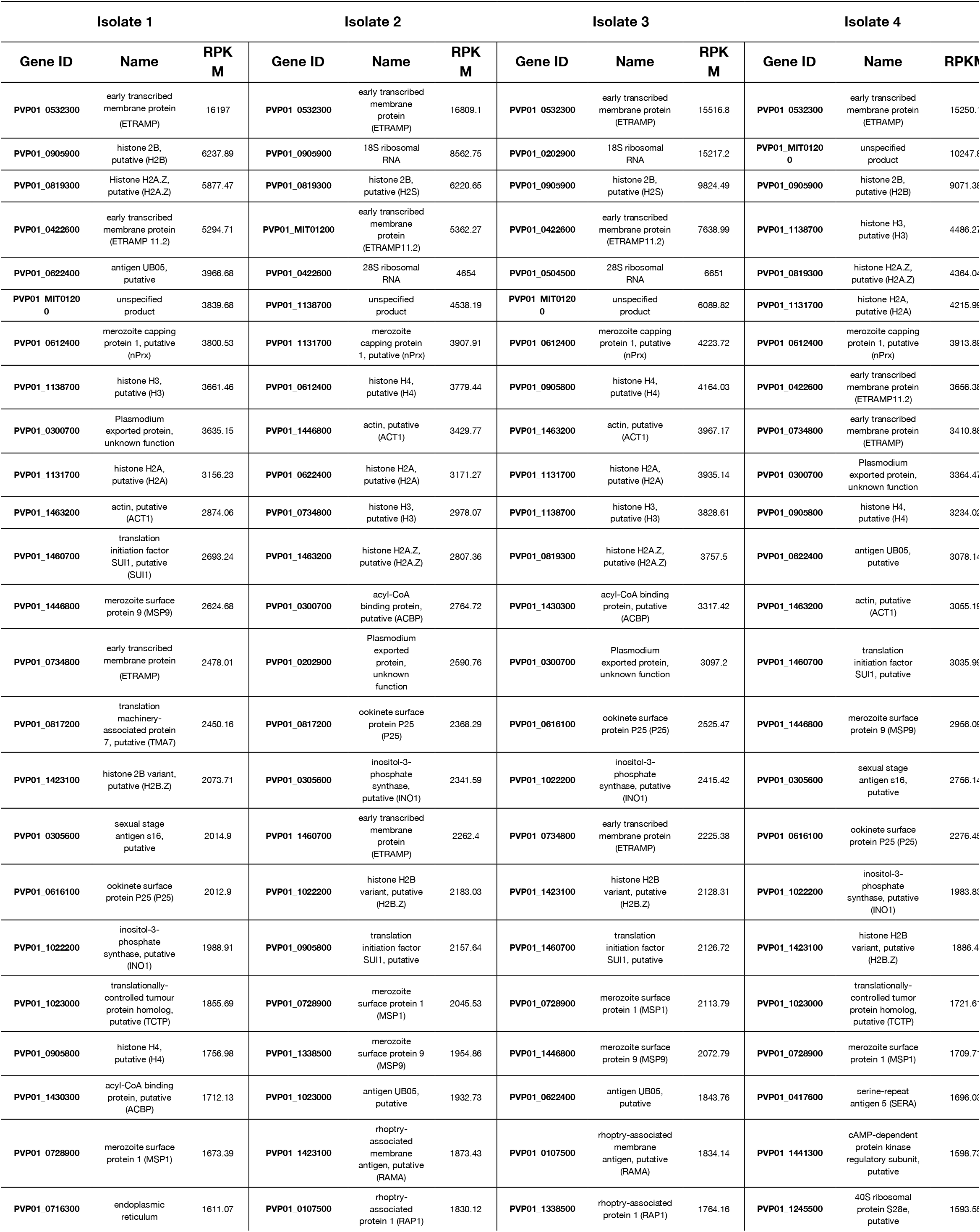
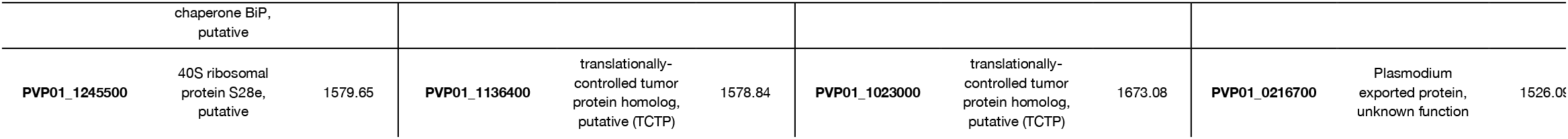
List of the 25 most expressed genes in each sample (RPKM).

We used two metrics of expression variability, coefficient of variation (*Cv*) and the index of dispersion (*D*), to identify genes that are variable between patient isolates. The intersection of the top 300 ranked genes derived using each metric were used to create a list of 86 genes that were differentially expressed between the isolates (**Supplemental Table 3.1**). Because many of the most variably expressed genes belong to multigene families known to have considerable sequence variation compared to the reference genome, we next evaluated the extent to which read mapping difficulties due to sequence variation were impacting read counts (and therefore downstream expression RPKM calculations). RPKM calculations leverage the number of reads that map to a gene across the entire length of coding sequence for each gene, while normalising for sequencing depth and gene length, giving reads per kilobase million mapped reads. In order to minimise the impact of read mapping drop out that would occur in some regions of a gene that have high sequence variation, we calculated RPKM values from reads only in regions of coding sequence where all isolates had at least five mapped reads, and stitched these regions together to re-calculate RPKM values for each gene (**Supplemental Table 3.2, Fig. S9**). This method, which allowed regions of considerable sequence divergence between the isolates to be ignored and hence true expression variation between each of the isolates to be assessed without the impact of sequence variation, resulted in a list of 104 differentially expressed genes (**Supplemental Table 3.2**). All of the 86 differentially expressed genes found in the analysis of the full coding regions were found in this second round of analysis of only conserved transcribed regions. Of the final list of 104 differentially expressed genes, 23 were not present in the Sal1 genome assembly, highlighting the importance of using the most complete annotation available.

A large proportion of the 104 genes shown to be differentially expressed between the four isolates occurred in several gene cluster hotspots, the largest of which are on chromosomes 5 and 10 (**Table 2**). Many differentially expressed genes belong to several multigene families implicated in immune evasion, antigenic variation and virulence, such as the *msp, phist, vir* and *trag* families (**Table 2**). Several other gene families involved in host cell recognition were also found to have differential expression profiles, specifically the reticulocyte binding proteins (RBPs) and tryptophan-rich proteins (TRAG) and duffy binding protein (DBP) (**Table 2**). There was no correlation between genes that had the highest levels of expression and those with highest variability between isolates, which was expected because as noted above, the top expressing genes were highly correlated between isolates. We suggest that these genes may be subject to epigenetic variation, as specific family members are differentially expressed across the four isolates. There is direct experimental evidence for epigenetic regulation multigene families in other *Plasmodium* species, such as the *pir* gene family ^27^ (which includes the *P. vivax vir* genes) and experiments with clonal parasites have shown that epigenetic regulation occurs in numerous gene families in *P. falciparum* ^28^.

**Table 2.** Differentially expressed genes belonging to clusters (in bold) or multigene families in Cambodian isolates (RPKM-adjusted for sequence variation).

| Gene ID | Name | Isolate 1<br>RPKM | Isolate 2<br>RPKM | Isolate 3<br>RPKM | Isolate 4<br>RPKM | C | D |
| --- | --- | --- | --- | --- | --- | --- | --- |
| PVP01_0000170 | tryptophan-rich protein (TRAG32) | 29.27 | 240.68 | 43.30 | 127.00 | 0.88 | 85.81 |
| PVP01_0004360 | PIR protein | 65.82 | 88.33 | 148.22 | 7.42 | 0.75 | 43.75 |
| PVP01_0004370 | PIR protein | 162.13 | 55.23 | 24.47 | 25.21 | 0.98 | 63.63 |
| **PVP01_0010200** | merozoite surface protein 3, putative | 1246.43 | 3.94 | 1.49 | 1.90 | 1.98 | 1234.29 |
| **PVP01_0010220** | merozoite surface protein 3, putative | 88.55 | 59.88 | 268.46 | 367.45 | 0.75 | 110.02 |
| PVP01_0122100 | PIR protein | 89.09 | 24.60 | 22.38 | 11.90 | 0.95 | 33.44 |
| PVP01_0201800 | PIR protein | 132.21 | 399.29 | 124.40 | 431.42 | 0.61 | 101.71 |
| PVP01_0405200 | Plasmodium exported protein (PHISTc) | 24.31 | 12.49 | 87.81 | 8.58 | 1.11 | 41.02 |
| PVP01_0417400 | serine-repeat antigen 4 (SERA) | 96.58 | 42.93 | 61.45 | 192.20 | 0.68 | 44.91 |
| PVP01_0423700 | PIR protein | 51.79 | 11.96 | 2.67 | 7.34 | 1.22 | 27.59 |
| **PVP01_0504000** | Plasmodium exported protein (PHIST) | 102.08 | 54.14 | 157.91 | 38.34 | 0.61 | 32.91 |
| **PVP01_0504100** | Plasmodium exported protein | 19.10 | 7.17 | 69.67 | 5.00 | 1.20 | 36.29 |
| **PVP01_0504400** | sporozoite invasion-associated protein 2, putative (SIAP2) | 56.42 | 17.14 | 120.93 | 8.91 | 1.01 | 51.38 |
| **PVP01_0504500** | 28S ribosomal RNA | 973.82 | 1698.50 | 9873.71 | 1206.48 | 1.25 | 5380.46 |
| **PVP01_0504700** | 18S ribosomal RNA | 80.00 | 202.88 | 963.80 | 84.93 | 1.27 | 541.09 |
| PVP01_0515900 | Plasmodium exported protein (PHIST) | 67.25 | 29.84 | 418.41 | 18.47 | 1.43 | 273.52 |
| PVP01_0516500 | Plasmodium exported protein (PHIST) | 15.97 | 9.14 | 57.87 | 5.37 | 1.10 | 26.64 |
| PVP01_0523200 | Plasmodium exported protein (PHIST) | 42.97 | 25.14 | 95.39 | 11.34 | 0.84 | 30.99 |
| PVP01_0524100 | Plasmodium exported protein (PHIST) | 33.26 | 13.50 | 83.18 | 14.15 | 0.91 | 29.77 |
| PVP01_0533700 | Plasmodium exported protein (PHIST) | 45.73 | 21.98 | 238.70 | 9.16 | 1.36 | 146.77 |
| PVP01_0534300 | reticulocyte binding protein 2c (RBP2c) | 62.67 | 41.64 | 177.40 | 14.22 | 0.97 | 69.57 |
| **PVP01_0601600** | Plasmodium exported protein (PHIST) | 24.48 | 15.11 | 74.78 | 5.13 | 1.04 | 32.09 |
| **PVP01_0601700** | Plasmodium exported protein (PHIST) | 54.34 | 30.64 | 161.54 | 15.87 | 1.00 | 66.20 |
| PVP01_0623800 | duffy binding protein (DBP) | 90.60 | 32.36 | 277.41 | 20.05 | 1.13 | 134.55 |
| PVP01_0700700 | tryptophan-rich protein (TRAG34) | 136.11 | 84.55 | 310.05 | 52.34 | 0.79 | 90.46 |
| PVP01_0701200 | reticulocyte binding protein 1a (RBP1a) | 70.02 | 31.49 | 152.90 | 20.69 | 0.87 | 52.25 |
| PVP01_0800700 | reticulocyte binding protein 2b (RBP2b) | 45.68 | 18.96 | 99.58 | 12.59 | 0.90 | 35.49 |
| PVP01_0808700 | Plasmodium exported protein (PHIST) | 19.84 | 6.29 | 121.89 | 10.69 | 1.39 | 76.52 |
| PVP01_0839300 | PIR protein | 68.46 | 7.34 | 5.55 | 13.42 | 1.27 | 38.08 |
| **PVP01_1031100** | merozoite surface protein 3 (MSP3G) | 52.07 | 67.67 | 196.89 | 43.37 | 0.80 | 57.55 |
| **PVP01_1031400** | merozoite surface protein 3 (MSP3.5) | 143.78 | 78.00 | 14.38 | 84.49 | 0.66 | 34.92 |
| **PVP01_1031500** | merozoite surface protein 3 (MSP3.3) | 208.61 | 185.68 | 94.54 | 835.10 | 1.03 | 348.59 |
| **PVP01_1031700** | merozoite surface protein 3 (MSP3.1) | 2606.85 | 1761.98 | 1889.06 | 197.99 | 0.63 | 637.85 |
| PVP01_1033800 | tryptophan-rich protein (TRAG17) | 402.99 | 155.61 | 853.86 | 195.32 | 0.80 | 255.09 |
| PVP01_1100400 | PIR protein | 50.33 | 167.19 | 25.26 | 55.55 | 0.85 | 53.45 |
| PVP01_1201400 | Plasmodium exported protein (PHIST) | 57.95 | 34.49 | 215.77 | 12.58 | 1.15 | 106.14 |
| **PVP01_1219800** | MSP7-like protein (MSP7.6) | 179.79 | 177.55 | 496.23 | 76.87 | 0.78 | 142.69 |
| **PVP01_1220200** | MSP7-like protein (MSP7.9) | 1032.53 | 2939.70 | 1457.97 | 322.91 | 0.77 | 849.01 |
| PVP01_1401800 | tryptophan-rich protein (TRAG21) | 34.54 | 22.27 | 118.27 | 6.70 | 1.10 | 54.72 |
| PVP01_1402400 | reticulocyte binding protein 2a (RBP2a) | 44.16 | 16.73 | 95.14 | 11.52 | 0.91 | 34.98 |

The sequence-variation adjusted RPKM analysis confirmed that the vast majority of highly-expressed genes in each isolate were conserved compared to the initial analysis of differential expression results, with only one of two genes changing in the top 25 highest expressed genes for each isolate, which would be expected to stay largely the same with the use of these strict parameters for mapping (**Supplemental Table 3.2**). For the genes with highest variability between isolates, nearly all the genes from the original analysis remained significantly variably expressed (81/86), with the exception of five genes including two *msps* (*msp3.9* and *MSP3.2*). *MSP3.9* and *MSP3.2* appear to be an example of sequence variation causing variable levels of mapping instead of true expression variation, as seen in **Fig. 3a** where mapping of the *msp3* locus shows individual isolates with reads mapping for some isolates but not others. For *msp3.9*, isolate 4 shows a distinct peak of expression in the middle of the transcript, where the other isolates drop off in expression likely due to sequence variation, and adjusted analysis corrects for these differences, finding that there is no longer a significant variation in expression for the four isolates after correction (**Fig. 3a**). Similarly, the *msp3.2* locus has a similar profile with high levels of expression for isolate 2 that is lost upon RPKM correction (**Fig. 3a**). Adjusted RPKM analysis additionally found 23 new genes to be highly variably expressed, including six *pir* genes and several putative exported proteins, many of them belonging to the *phist* superfamily which were the most represented of all the gene families, as well as reticulocyte binding protein 2c (PVP01_0534300) (**Supplemental Table 3.2**).

**Figure 3:**
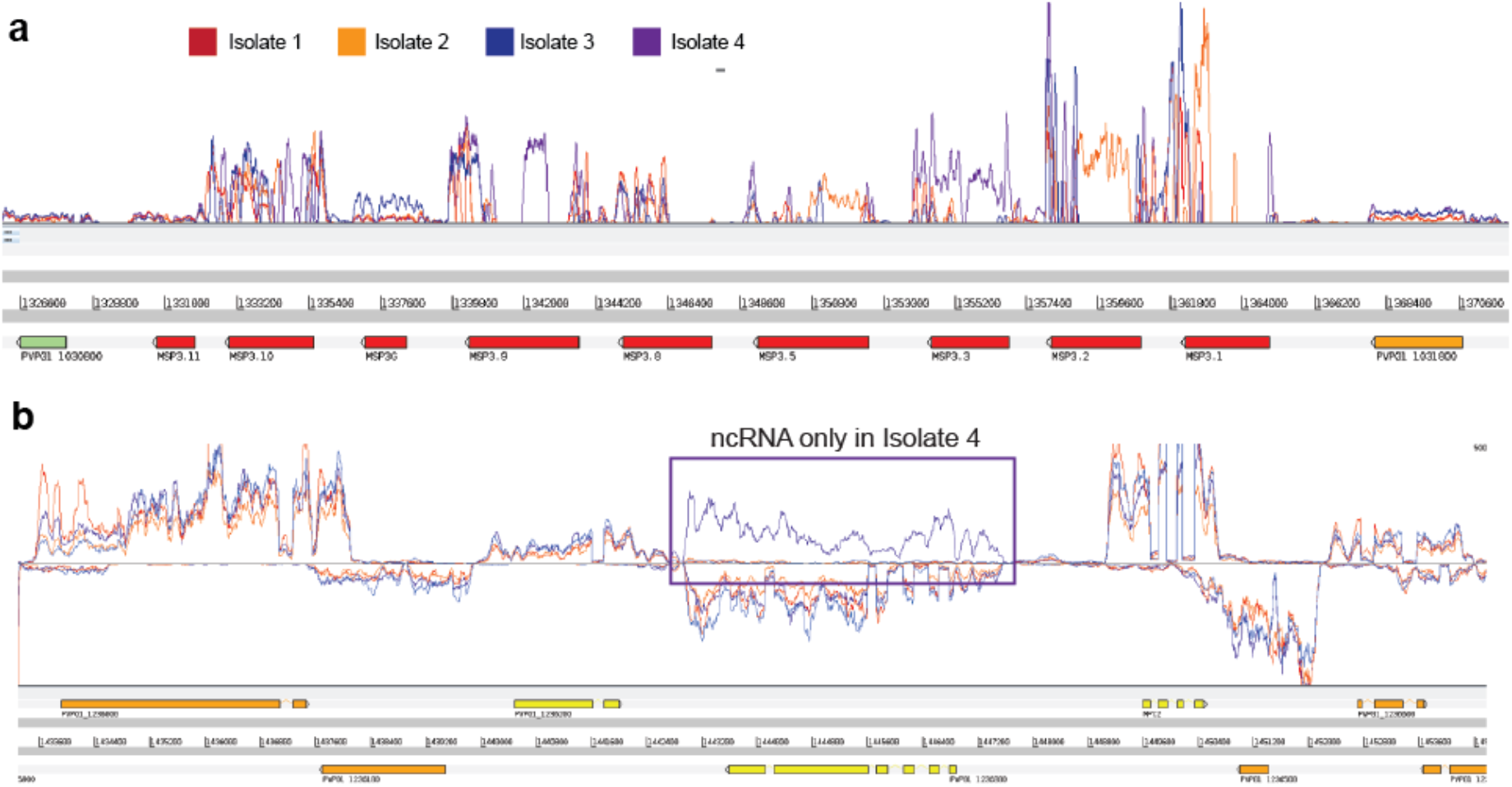
Variation in RNA-seq data between the four patient isolates. a. RNA-seq coverage data (lines in top panel) for the region on PvP01 chromosome 10, which contains the sequences of multiple MSP3 genes (shown in red on the lower panel). Levels of transcripts detected for each of the genes in this locus vary between each of the patient isolate (see key in figure to identify traces). Drops in coverage within an annotated gene model are likely to represent sequence divergence in the patient isolates from the reference genome. b. RNA-seq coverage data (lines in top panel) for a region of PvP01 chromosome 10. The four patient isolates show very similar transcript levels for most of the protein coding genes in this region. However, an unannotated ncRNA (highlighted by a box) was only found in isolate 4, opposite to the gene PVP01_1236300.

## Discussion

The data presented in this study enabled us to examine the extent of the *P. vivax* schizont transcriptome in samples derived from multiple patient isolates. We were able to identify transcripts originating from 75% of annotated protein coding genes, and to infer transcription from 78% of the genome sequence.

Our analysis has increased the amount of *P. vivax* genome sequence that can be annotated as encoding 5’ and 3’ UTRs. The median length of the 5’ UTRs (n =4155) is our data set was 815 nt, and was 685 nt for 3’ UTRs (n=4,091). In the first *P. vivax* RNA-seq study ^14^, the median length of 5’ UTRs was 295 nt (n=3,633), and the median length of the 3’ UTRs was 203 nt (n=3,967). In a more recent *P. vivax* RNA-seq study^15^, the median lengths of the was 754 nt for 5’ UTRs and 785 nt for the 3’ UTRs (n =3,230); the lack of a difference in relative lengths of the 5’ and 3’ UTRs is likely to be a feature of the computational approach used in that work, which did not consider each UTR independently. The improved annotation of UTRs in this data was enabled by a more even coverage of AT-rich regions of the genome compared to other studies, in tandem with a computational approach developed specifically for *Plasmodium* genomes. This custom approach makes use of blocks of continuous RNA-seq coverage to identify UTRs, so is less likely to merge fragments of RNA-seq coverage from independent gene models than approaches that are built for genomes where gene models are further apart (such as mammalian genomes) Future studies comparing similarities in UTRs between and within *Plasmodium* species may be able to identify RNA-binding motifs relevant to gene regulation, though it should be noted that the AT-rich nature of these sequences restricts application of existing motifs for predicting RNA structure. The extended UTR sequences enabled identification of hundreds of overlapping transcripts pairs, even though all of the UTRs were called within the same life cycle stage. This observation suggests levels of complexity in gene regulation in *P. vivax* that are not yet fully understood, as only one DNA strand can be transcribed at any given time. Future single cell RNA-seq studies may be able to clarify this, but will require understanding of UTR boundaries to interpret sparse and noisy data.

Detection of previously undescribed features of the *P. vivax* transcriptome will enable a more nuanced understanding of gene regulation in this species. The detection of the presence of TSS-associated RNAs in *P. vivax* reveals a new class of RNAs in *P. vivax*, however it remains to be proved whether these transcripts play a direct regulatory role (such as binding proteins near active TSS), or are a byproduct of transcription of mRNAs that has no further function. The extended catalogue of splice sites in *P. vivax* will help to better understand gene regulation within the species. We note that other studies have identified splice sites^14,15^, but this study is the first to identify the presence of exitrons in *P. vivax*. Exitrons were first described in humans and plants^26^, and can be used to increase proteome plasticity as well as to regulate transcript levels through alternative splicing, including transcript degradation by nonsense-mediated decay. Exitrons were recently described in *P. falciparum*^*23*^, and may prove to be an important component of gene regulation across the *Plasmodium* genus.

The use of a Percoll gradient to generate matched-stage schizont samples from multiple patient isolates has enabled an analysis of genes that are differentially expressed between the parasites isolated from each patient. Most genes have highly similar patterns of expression between the patient isolates, enabling differences in staging to be removed from the analysis. This enables focus on the genes that are truly variable between patient isolates. Of particular interest was how schizont transcription from patient infections are different to one another, as the transcriptional heterogeneity could have profound impacts on the host immune response and perhaps even the invasion phenotypes of the parasites. Prior studies have implicated that high levels of invasion related proteins like Duffy binding protein (DBP1) could enable parasites to invade duffy negative erythrocytes, and *P. vivax* may additionally have alternative invasion pathways independent of DBP1^29–31^. Recent studies of SalI parasites infecting *Aotus* and *Saimiri* monkeys slooked at potentially unique invasion pathways and implicated the differential expression of the tryptophan-rich protein family in mediating this process^18^. No clear pattern of differential expression was shown in the four patient isolates studied, however the expression heterogeneity of these erythrocyte-binding proteins in this study and several others warrants further investigation about the involvement of these in host recognition.

Many of the genes most variably expressed between the isolates appear to cluster in physical space along regions of the chromosomes (**Table 2**), such as multiple members of the MSP3 family, which are known to be under heavy diversifying selection^32,33^, as shown in **Fig. 3**. Levels of higher and lower transcription within each cluster of variably expressed genes suggest that the parasites are making choices about which version of the gene to express within a patient. Epigenetic regulation of multigene families has been studied in *P. falciparum*, with mechanisms capable of selecting specific members of gene families thought to be observed across the genus^34^. It is likely that *P. vivax* parasites are capable of selecting particular members of gene families to be expressed in response to the host, which is supported by recent evidence describing the response of *P. vivax* parasites to different primate hosts^18^. Our evidence of putative spatial relationship between genes which are variably expressed suggests that chromatin structure and accessibility may be important in this class of gene regulation, as transcriptional heterogeneity has been implicated to be of considerable importance in parasite survival to changing host conditions^28,35^. In the case of the *msp3* locus, the paralogs are likely examples of functional redundancy that acts to ramp up antigenic diversity, enhancing the ability to evade the immune system during invasion^32^. For other multigene families, such as the *vir* genes, are subtelomeric genes involved in the establishment of chronic infections, and have been shown to be regulated independently of the cell cycle. Clonally-variant expression seen in *vir* genes has been established previously, and indeed we found that all the *vir* genes differentially expressed in the current study had evidence varying amounts of expression, and unsurprisingly, had no obvious on/off transcriptional pattern^36^. Overall, this bet-hedging strategy is an effective one where stochastic changes in expression levels between individual parasites spreads out the immunogenic risk among the population, allowing selection on existing variation and survival of the fittest among them. Future studies of *P. vivax* transcriptomes should consider this layer of heterogeneity.

## Methods

### Field isolate collection and schizont enrichment

Samples were selected for ex vivo culture from *P. vivax* malaria patients presenting to Sampov Meas Referral Hospital, Cambodia. These patients had not taken antimalarials within 1 month of sample collection and had parasitemia of <0.1%. Written informed consent was obtained from each donor. Prior approval of the clinical study protocols were obtained from the National Ethics Committee for Human Research (Clinical Trials.gov Identifier: NTC00663546) in Cambodia, and by the Institutional Review Board, NIAID, NIH. The clinical trial was conducted and data were generated and biological specimens were collected, and reported in compliance with the study protocol (approved by the Institutional Review Board, NIAID, NIH and the National Ethics Committee for Human Research in Cambodia), International Conference on Harmonisation Good Clinical Practice (ICH GCP), and Good Laboratory Practices (ICH GLP). Sample processing in the field was undertaken as previously described (Russell et al., 2011). Briefly, four *P. vivax* isolates were processed (PV0417-3, PV0563, PV0565, PV0568). Patient blood samples (16 ml for ages <18 years and 32 ml for ≥18 years) were collected by venipuncture into sodium heparin vacutainers, then were centrifuged (2000 rpm for 5 min) and plasma was removed. Samples were diluted with PBS (up to 64 ml, mixed by inverting tube), depleted of white blood cells and platelets (pass over autoclaved, pre-wet CF11 columns packed to the 5.5-ml mark in a 10-ml syringe, and collect flow through), washed twice (centrifuged at 2000 rpm for 5 min, washed with 1x PBS, repeated 1 time), and placed into *ex vivo* culture conditions (resuspended with packed cells at a 10% hematocrit in modified McCoys 5A complete media with 25% AB serum, and cultured at 37°C, 5% CO_2_) until the parasites matured to schizonts as identified by Giemsa stained microscopy. After maturation, cultures were pelleted (centrifuge at 2000 rpm for 5 min), the supernatant was removed, and cells were resuspended (to 50% hematocrit in 1x PBS). To prevent rosetting, cells were treated with trypsin (7.5 ml of 500 mg/l Trypsin-Versene) and incubated at 37°C for 15 min. To stop digestion, samples were diluted (add 2x volume of 1x PBS) and centrifuged (2000 rpm for 5 min). The recovered pellet was then incubated with AB serum (6 ml for 5 min at RT), diluted with PBS (up to 30 ml) and separated on a 45% isotonic Percoll gradient (5 ml of suspension overlaid on six 15-ml tubes containing 5 ml 45% isotonic Percoll each). The suspensions were centrifuged (1200 RPM for 15 min) and the fine band of concentrated schizonts on the Percoll interface was removed, centrifuged (2000 rpm for 5 min), and resuspended (5 ml of 1x PBS and counted via hemocytometer). The remaining sample was pelleted, mixed with up to 10 volume of RNAlater (1 ml) and divided into 2 cryovials (500 μl each).

### RNA extraction and assessment of RNA purity by qPCR

RNA was isolated using the RiboPure Blood kit (Ambion) according to manufacturer instructions and subjected to 2 rounds of DNA digestion using the DNA-free kit (Ambion) according to manufacturer instructions. Samples were analysed with a Bioanalyzer 2100 (Agilent Technologies, Inc.) using an RNA Nano Chip to test quantity and quality. To test for genomic DNA contamination, 1 μl of RNA solution was used to make cDNA using the High Capacity cDNA Reverse Transcription Kit (Applied Biosystems) according to manufacturer instructions. Using both RNA and cDNA samples, a region surrounding an intron in the PvDBP gene PVX_110810 (5’:AAACCGCTCTTTATTTGTTCTCC, 3’:TTCCTCACTTCTTCTTTCATT) was amplified by PCR. Reaction volumes were as follows: 2.5 μl buffer, 2 μl dNTP mix (10 μM), 0.1 μl Platinum Pfx DNA Polymerase (Invitrogen), 16.9 μl water, 1 μl of each primer (10 μM), 1 μl cDNA or extracted RNA. Thermocycler conditions were as follows: incubation at 95°C for 15 min, 35 cycles consisting of denaturation at 95°C for 40 seconds, annealing at 55°C for 40 seconds, elongation at 68°C for 1 min, followed by a final extension at 65°C for 5 min.

### RNA-seq libraries

A modified RNA-seq protocol (“DAFT-seq”^23^) was used to capture schizont transcriptomes generated in this study. Briefly, polyA+ RNA (mRNA) was selected using magnetic oligo-d(T) beads. The polyA+ RNA (mostly mRNA) was fragmented using a Covaris AFA sonicator using the following settings: Duty cycle 10%, Intensity 5, Cycles per burst 200, Time 60s, then was precipitated using ethanol. Reverse transcription using Superscript II (Life) was primed using random primers in a 20 uL reaction, the RNA-DNA hybrid was cleaned using 1.8x reaction volume with Agencourt RNAClean XP, then second strand cDNA synthesis with DNA pol I and RNase H, and included dUTP. A “with-bead” protocol was used for dA-tailing, end repair and adapter ligation (NEB) using “PCR-free” barcoded sequencing adaptors (Bioo Scientific, similar to Kozarewa *et al.* ^37^. After 2 rounds of SPRI cleanup the libraries were eluted in EB buffer and USER enzyme mix (NEB) was used to digest the second strand cDNA, generating directional libraries. The libraries were subjected to 4 cycles of PCR using the 2x KAPA HIFI HS Master mix. Reaction conditions were as follows: 95°C for 5 min followed by 4 cycles of 95°C for 20 s, 60°C for 15 s, 72°C for 60, and a final extension of 72°C for 5 min, then cleaned using 1.0x reaction volume with Agencourt RNAClean XP. The libraries were quantified by qPCR and sequenced on an Illumina HiSeq2000 using 100 bp paired end reads.

### Sequence mapping and quality control

The PvP01 reference genome^9^ and the January 2019 GeneDB annotation release were used for all analysis. Sequencing data from these samples has been made publicly available in the European Nucleotide Archive (ENA) under Study ID PRJEB32240. TopHat2^38^ was used to map reads with directional parameters and a maximum intron size of 5,000 nt. RNA-seq data was visualised using the Artemis genome browser^39,40^.

### Identification of transcriptome features

Custom Perl scripts were used to calculate RPKM values, which used the BEDTools suite^41^. A custom approach described previously^23^ was used to detect new RNA sequences, including UTRs and ncRNAs, which also used the BEDTools suite. Spiced reads were identified using the CIGAR string in the DAFT-seq BAM files. Comparison with existing annotation allowed the detection of known spice sites. Splice sites in UTRs and ncRNAs were found by identifying overlapping spliced reads. Exitrons were found by identifying spliced reads mapping within protein-coding exons. Detection of each splice site required at least 5 reads. Code is available online at http://github.com/LiaChappell/DAFT-seq

### Comparisons to other data sets

For comparison to microarray data sets, one-to-one orthologues were selected for comparison between time courses using Sal1^12^ and for with *P. falciparum* ^*24*^. Pearson correlation was calculated for orthologous pairs of genes present in both data sets. The corrplot R package was used to visualise the data.

### Comparative analysis of isolates

A threshold of 5 RPKM (reads per kilobase per million mapped reads, a normalised measure of gene expression) was used to define detection of expression. In order to compare relative levels of expression between the individual isolates to look for expression variability, two metrics were calculated for each gene, the coefficient of variation (*Cv*) which is a ratio of standard deviation (*σ*)of RPKMs from the four isolates to the mean (*μ*) RPKM:

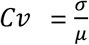

and the index of dispersion (*D*), which gives an indication of the spread of expression results, represented as:

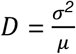

The intersection of the top ranked genes from each method were used to create a list of 86 differentially expressed genes between the isolates. Gene ontology (GO) terms and 1:1 *P. falciparum* homologs (29/86) were identified for differentially expressed genes using PlasmoDB^42^, and genes identified as part of the *P. falciparum* invadome were also evaluated against our dataset^43^.

To be able to differentiate between variable levels of expression and variable levels of mapping (due to sequence variation in the isolates), we performed an additional round of analysis considering only regions of coding sequence where reads were mapped in all of the isolates. We examined blocks of continuous coverage where at least 5 reads were mapped, breaking coding sequences into multiple blocks that could be effectively considered as exons in downstream analysis of differential expression (see **Fig. S7**).

### Data availability

Sequencing data from these samples has been made publicly available in the European Nucleotide Archive (ENA) under Study ID PRJEB32240.

## Acknowledgements

The authors would like to thank Thomas D. Otto for help regarding the PvP01 reference genome, Liam Prestwood for help with transport of samples and Mandy Sanders for help with sequencing submissions. The authors would also like to thank the staff of WSI Bespoke DNA pipelines team for their contribution. Funding support was provided by National Institutes of Health/NIAID (R01AI137154), the Intramural Program of the National Institute of Allergy and Infectious Diseases (National Institutes of Health) and the Wellcome Trust (206194/Z/17/Z).

## Author contributions

S.V.S. and L.C. performed data analysis, interpreted results and wrote the manuscript. J.B.H. purified the schizonts and extracted the RNA. J.B.H. and L.C. performed the RNA-seq library construction. C.A. and S.S. collected samples in field sites and prepared the blood samples in RNAlater. U.C.B. updated annotation of the gene models using the RNA-seq data. M.B., R.M.F. and J.C.R. contributed to manuscript writing and interpretation of results.

