## supplemental figures for "Analysis of *Plasmodium vivax* schizont transcriptomes from field isolates reveals heterogeneity of expression of genes involved in host-parasite interactions"

**Siegel et al. 2020. Supplemental figures S1-S9**

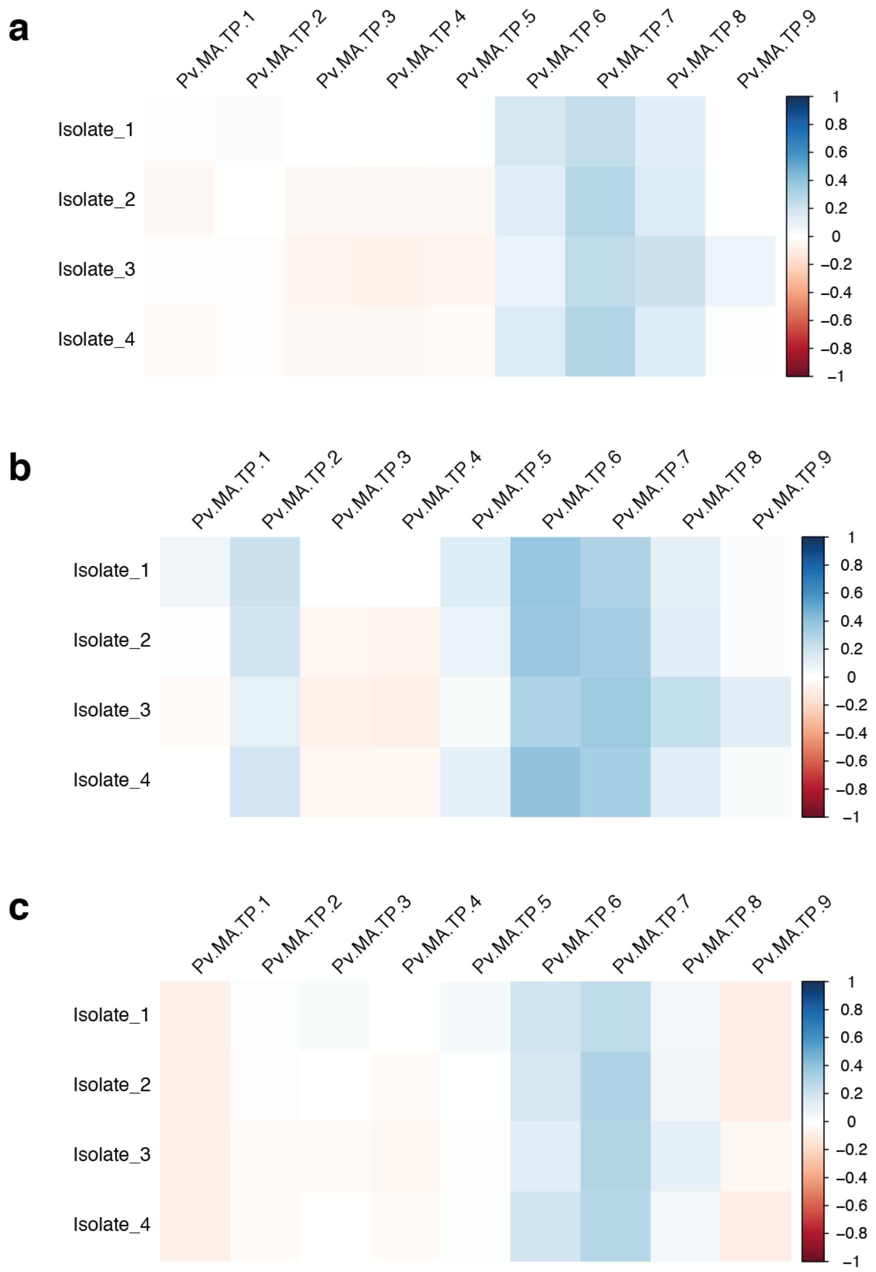


**Figure S1: Comparison of patient isolate RNA-seq to a *P. vivax* array blood stage array time course**

The expression levels of genes with one-to-one orthologues were compared between the RNA-seq data from the four patient isolates and microarray data from three IDC time courses from patient samples (Bozdech *et al*., 2008). The RPKM values from the RNA-seq data were compared to the fold changes in expression in the time course, where values were available for both orthologues (missing values were ignored). Despite different properties of the dynamic range in the data sets, it is clear that the isolate data mostly strongly correlates with the microarray data points towards the end of the time course, which correspond to schizonts. a. Microarray time course SD1. b. Microarray time course SD2. c. Microarray time course SD3.

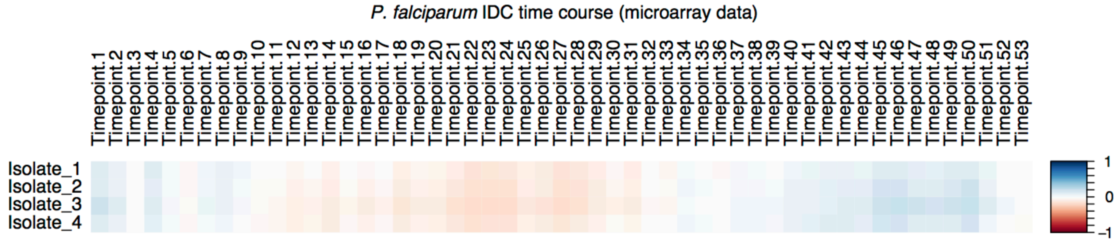


**Figure S2: Comparison of patient isolate RNA-seq to a *P. falciparum 3D7* IDC array time course**

The expression levels of genes with one-to-one orthologues were compared between the RNA-seq data from the four patient isolates and microarray data from a highly synchronised and densely sampled *P. falciparum* 3D7 IDC time course (Llinas *et al.*, 2006). The RPKM values from the RNA-seq data were compared to the fold changes in expression in the time course, where values were available for both orthologues (missing values were ignored). Despite different properties of the dynamic range in the data sets, it is clear that the isolate data mostly strongly correlates with the microarray data points towards the end of the time course, which correspond to schizonts.

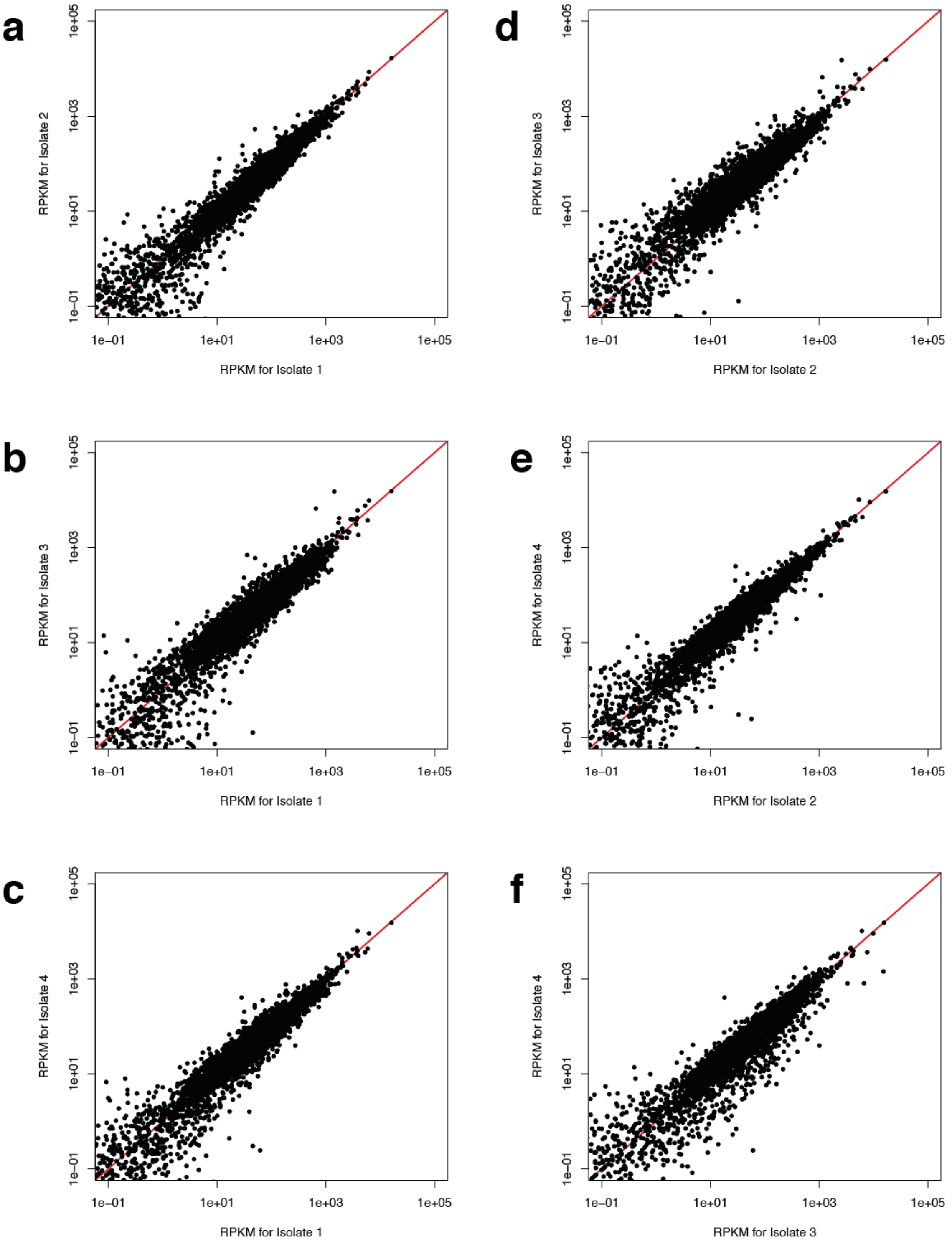


**Figure S3: Correlation plots for each pair of patient isolate RNA-seq samples**

RPKM values were compared for pairs of patient isolates, with data shown on a log scale. The red line indicates the line X=Y, with most points lying near this line, indicating that the patient isolates are very similar. a. Isolate 2 vs Isolate 1. b. Isolate 3 vs Isolate 2. c. Isolate 4 vs Isolate 1. d. Isolate 3 vs Isolate 2. e. Isolate 4 vs Isolate 2. f. Isolate 4 vs Isolate 3.

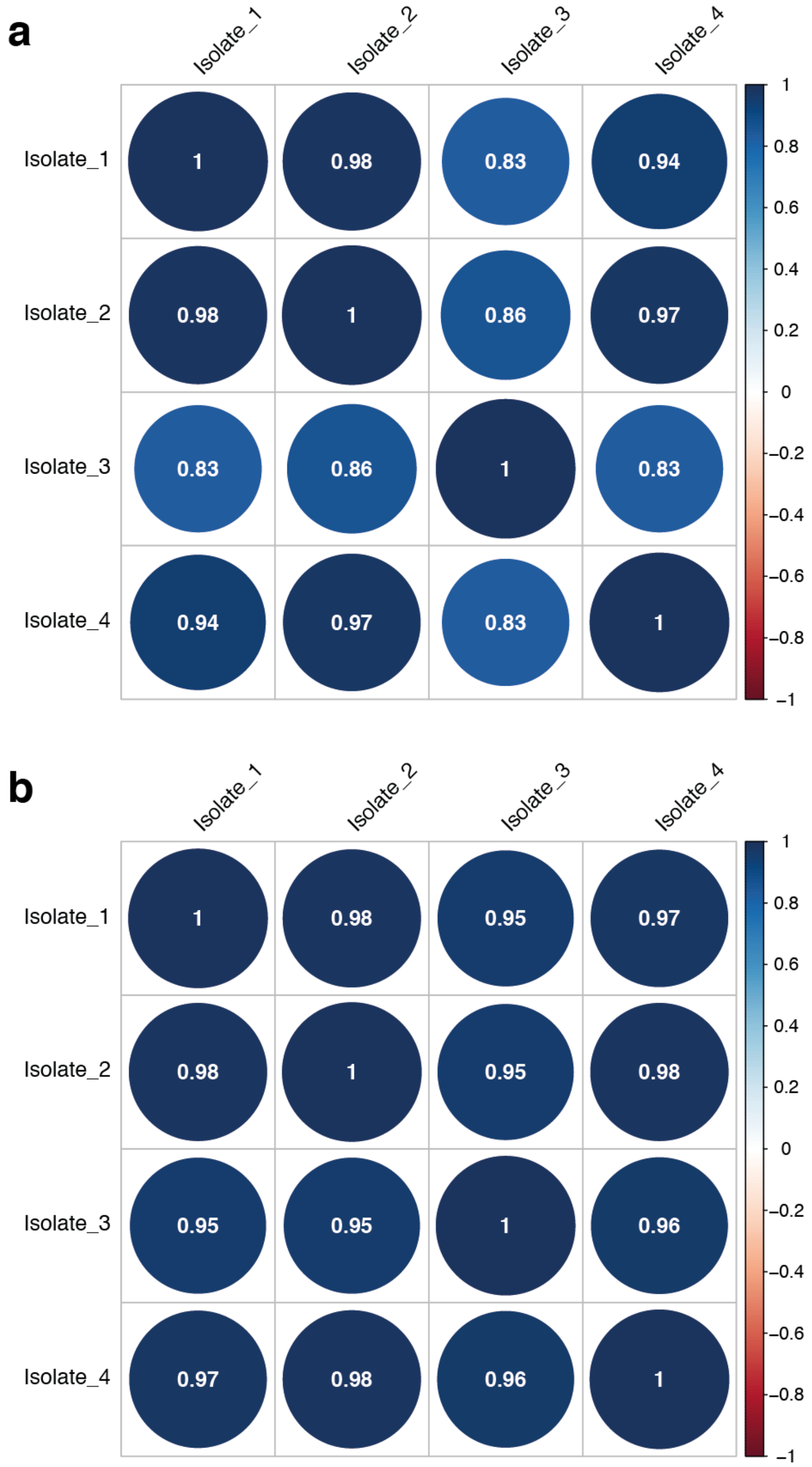


**Figure S4: Correlation values for each pair of patient isolate RNA-seq samples**

a. RPKM values were compared for each pair of isolates, using all of the gene IDs present in the PVP01 genome.
b. RPKM values were compared for each pair of isolates, using only gene IDs which were present in the Sal1 genome.

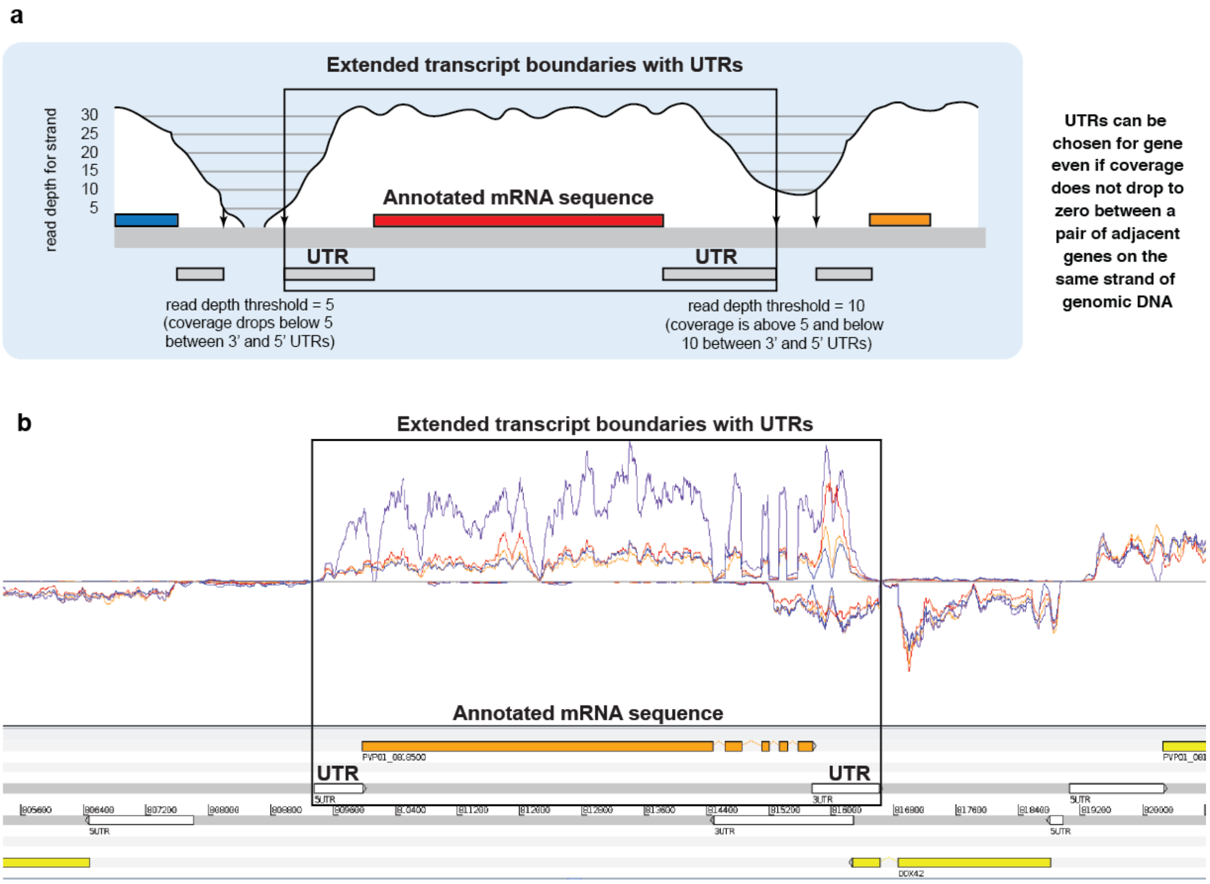


**Figure S5: Overview of UTR calling pipeline applied to *P. vivax* RNA-seq data**

a. Schematic view of UTR caller. More details can be seen in the supplementary material of the manuscript that first described the DAFT-seq approach (Chappell *et al.*).
b. Example of 5' and 3' UTRs called in this study extending the boundaries of a *P. vivax* gene model (PVP01_0818500).

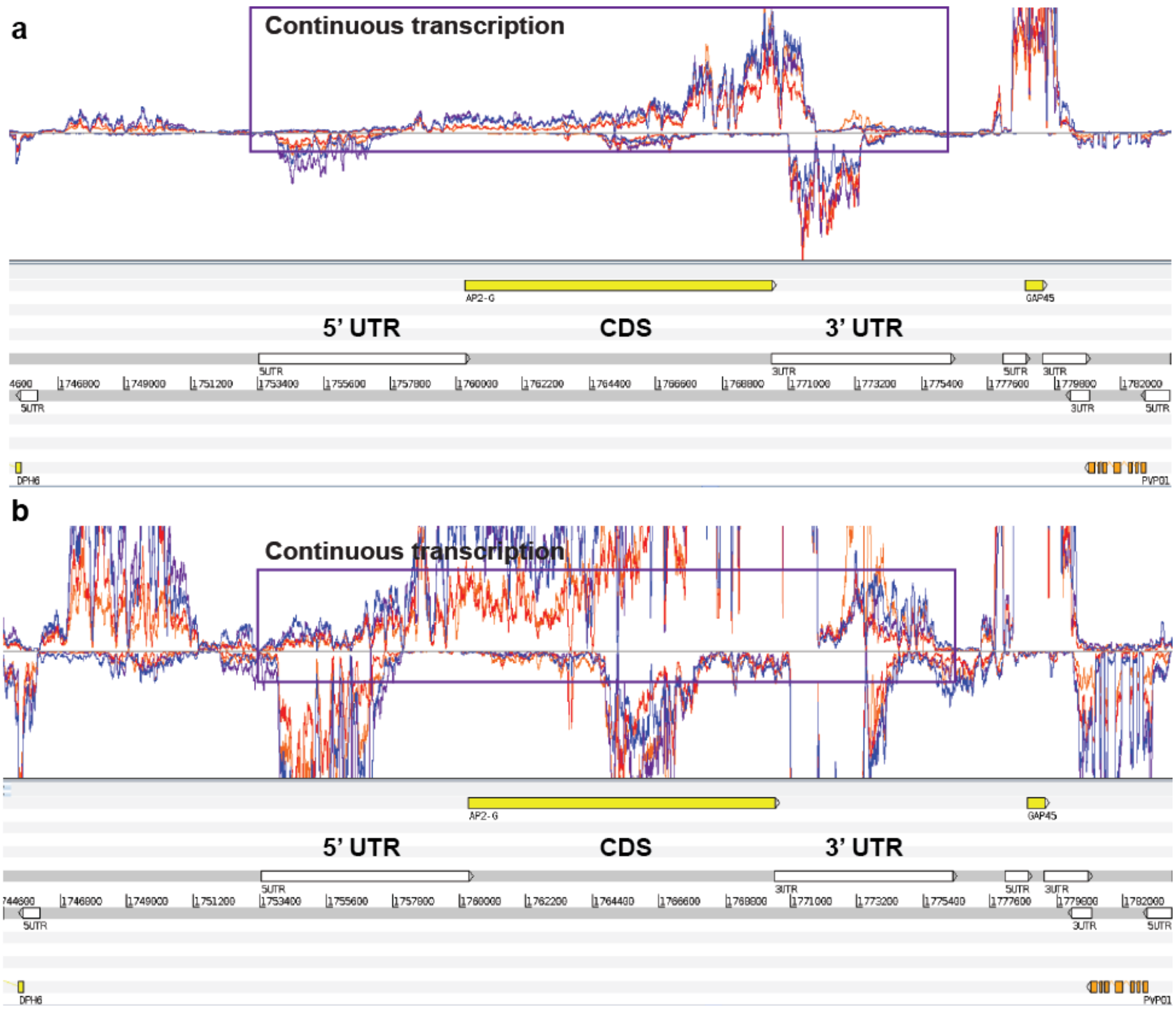


**Figure S6: The longest 5' and 3' UTRs belong to the gene AP2-G (PVP01_1440800)**

a. The colored traces represent RNA-seq coverage from each of the four patient isolates, which show very similar levels of coverage in this region. The box shows the detected boundaries of continuous RNA-seq coverage that overlap the AP2-G gene, showing all of the RNA-seq data.
b. A closer view (with "cropped" coverage) shows more clearly the boundaries of continuous coverage that overlap the the AP2-G gene. Coordinates of the boxes are the same in both panels.

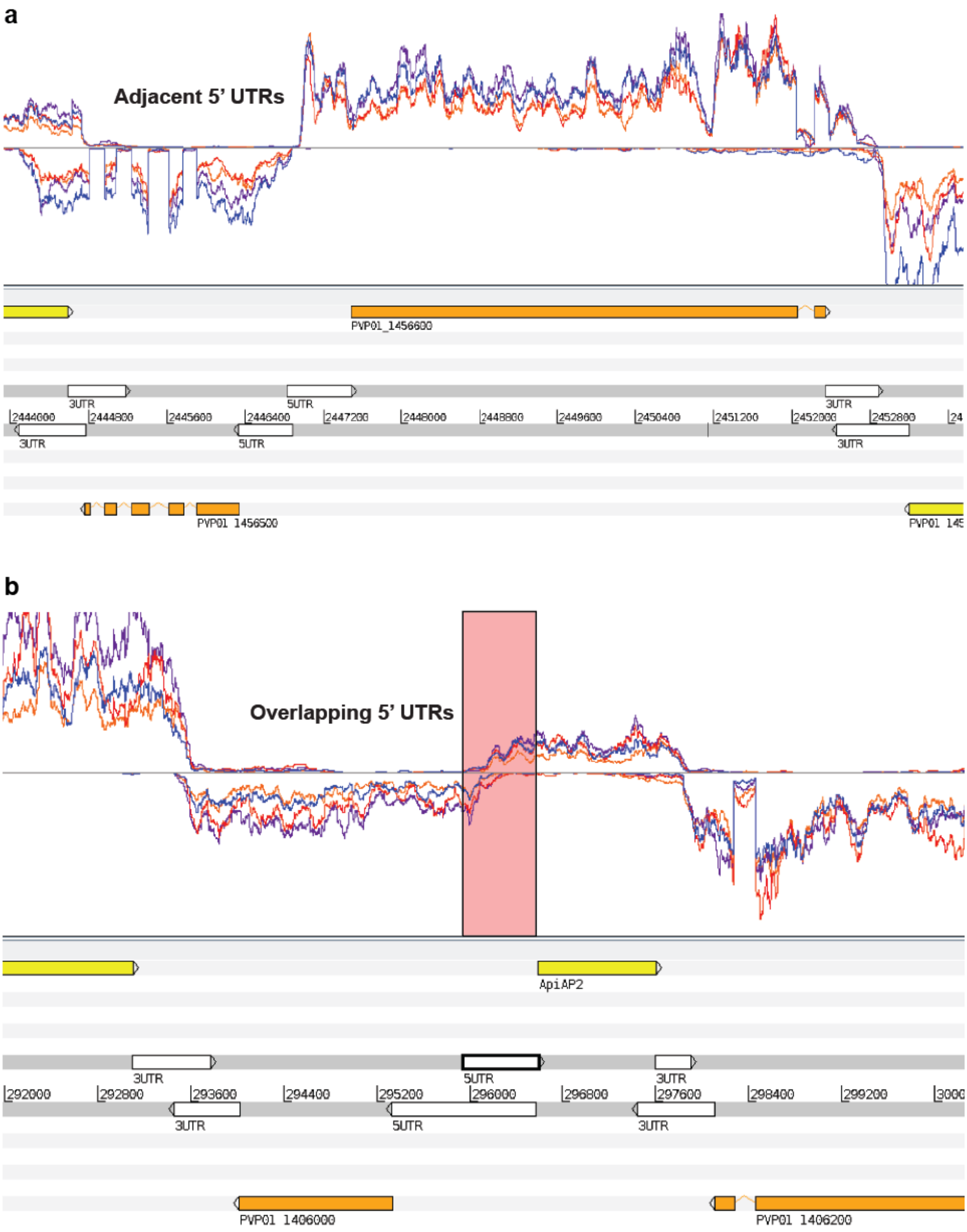
a
Adjacent 5' UTRs
PVP01_1456600
3UTR
5UTR
3UTR
2444000
2444800
2445600
2446400
2447200
2448000
2448800
2449600
2450400
2451200
2452000
2452800
3UTR
5UTR
3UTR
PVP01 1456500
b
Overlapping 5' UTRs
ApiAP2
3UTR
5UTR
3UTR
292000
292800
293600
294400
295200
296000
296800
297600
298400
299200
3UTR
5UTR
3UTR
PVP01 1406000
PVP01 1406200

**Figure S7: Adjacent and overlapping 5’ UTRs found in *P. vivax* schizonts**

a. The 5’ UTRs for the gene pair PVP01_1456500 and PVP01_1456500 are closely adjacent, and are likely to share a single bidirectional promoter.
b. The 5’ UTRs for the gene pair PVP01_1406000 and PVP01_1406100 (a gene encoding a protein with an ApiAP2 domain) significantly overlap, even though RNA was only collected from a defined life stage. A bidirectional promoter is likely to regulate this gene pair.

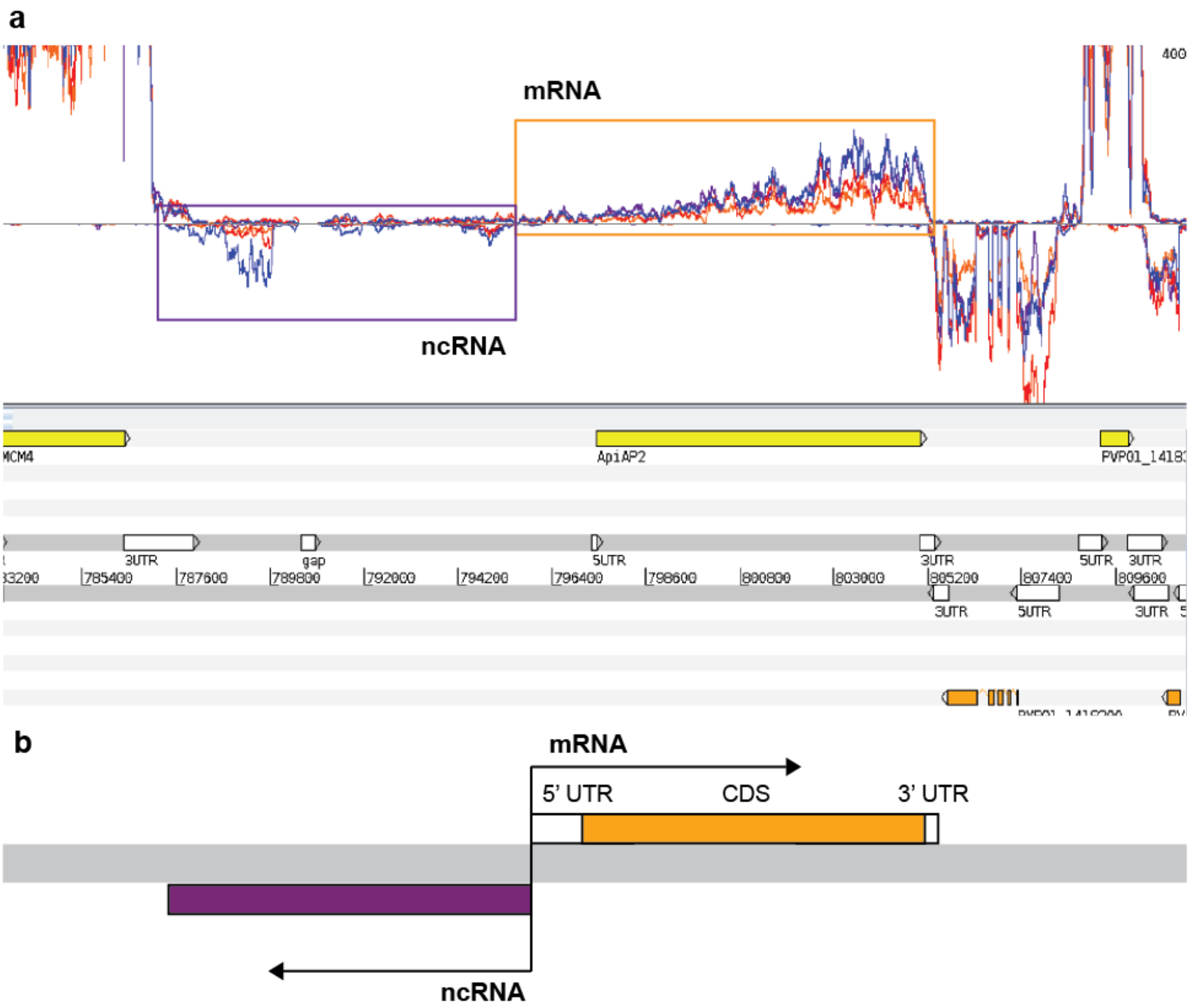


**Figure S8: TSS-associated RNAs are ncRNAs found in an antisense orientation to the 5' end of mRNAs, such as the example shown upstream of the gene AP2-G3 (PVP01_1418100)**

a. Boxes highlight RNA-seq coverage for the mRNA transcribed from the AP2-G3 gene (orange box on top panel) and a ncRNA in an antisense orientation to the 5' end of the gene (purple box). The pair of transcripts are likely to originate from a single bidirectional promoter.
b. Schematic view of the relative orientations of an mRNA and a TSS-associated RNA sharing the same bidirectional promoter.

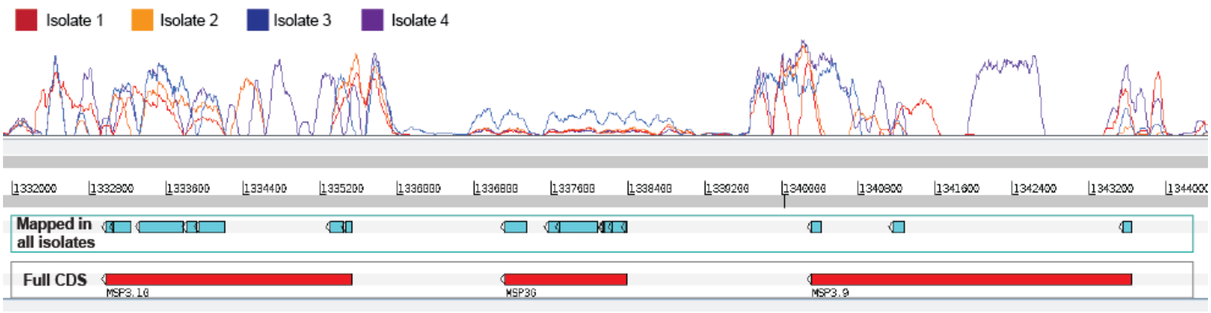


**Figure S9: Use of stricter parameters to analyse mapping in variable gene families**

For most genes, the mapping of RNA-seq coverage was very similar for each of the four isolates, indicating a comparable level of match to the reference genome. This can be observed in the other figures presented in this manuscript. However for some of the variable gene families, such as the MSP3 family (three members are shown in the figure above), the evenness of mapping along the length of the mRNA sequence is highly variable between the isolates. This suggests that there is variability in how similar the copies of these genes in each isolate are to the reference genome.

To be able to differentiate between variable levels of expression and variable levels of mapping (due to sequence variation in the isolates), we performed an additional round of analysis considering only regions of coding sequence where reads were mapped in all of the isolates. We examined blocks of continuous coverage where at least 5 reads were mapped, breaking coding sequences into multiple blocks that could be effectively considered as exons in downstream analysis of differential expression.

The figure above shows RNA-seq coverage (top panel) for the four isolates for three genes of the MSP3 family (left to right: MSP3.10, MSP3G, MSP3.9). In the lower part of the figure, the blue boxes show regions of the coding sequence that have mapped reads in all four isolates, while the red boxes at the bottom of the figure show the complete coding sequences annotated in the reference genome.
